# Relationship between serum bilirubin levels, urinary biopyrrin levels, and retinopathy in patients with diabetes

**DOI:** 10.1101/2020.11.23.393884

**Authors:** Kana Kudo, Tomoaki Inoue, Noriyuki Sonoda, Yoshihiro Ogawa, Toyoshi Inoguchi

## Abstract

**Aims/Introduction:** Previous reports have indicated that serum bilirubin levels may be associated with diabetic retinopathy. However, the detailed mechanism is not fully understood. In this study, we evaluated the relationship between the severity of diabetic retinopathy and various factors including bilirubin levels and factors influencing bilirubin metabolism.

**Methods:** The study participants consisted of 94 consecutive patients with diabetes mellitus admitted to Kyushu University Hospital from April 2011 to July 2012. The patients were classified into three groups: no retinopathy (NDR), simple retinopathy (SDR), and pre-proliferative or proliferative retinopathy (PDR). The relationship between the severity of retinopathy and various factors was evaluated using univariate and logistic regression analyses. In addition, multivariate regression analysis was performed to evaluate the significant determinants for bilirubin levels.

**Results:** In univariate analysis, a significant difference was found among NDR, SDR and PDR in bilirubin levels, duration of diabetes, systolic blood pressure, and macroalbuminuria. Logistic regression analysis showed that PDR was significantly associated with bilirubin levels, duration of diabetes, and systolic blood pressure (OR 0.737, 95% CI 0.570–0.952, P=0.012; OR 1.085, 95% CI 1.024–1.149, P=0.006; OR 1.036, 95% CI 1.011–1.062, P=0.005, respectively). In turn, multivariate regression analysis showed that bilirubin levels were negatively associated with high-sensitivity Creactive protein levels and PDR, but positively correlated with urinary biopyrrin levels, oxidized metabolites of bilirubin.

**Conclusions:** PDR was negatively associated with bilirubin levels. This negative association may be due to a decreased production of bilirubin rather than its increased consumption considering the positive association between bilirubin and biopyrrin levels.

## INTRODUCTION

Diabetic retinopathy is a common microvascular complication of diabetes, and a leading cause of visual impairment and blindness, especially in working-age adults (1, 2). Various metabolic abnormalities induced by chronic hyperglycemia have been postulated as causative factors for diabetic retinopathy (3). Among them, oxidative stress has gained increasing attention in recent years (4–7)). Oxidative stress is a pathological state and reflects an imbalance between reactive oxygen species (ROS) production and antioxidant capacity. Various reports have implicated both increased levels of ROS production and/or decreased antioxidant capacity in diabetic retina (5–7).

Bilirubin is mainly generated from heme degradation by heme oxygenase-1 (HO-1). Although bilirubin was once considered to be a toxic waste product of heme catabolism in human, it has recently emerged as an important strong antioxidant and cytoprotectant (8). An increasing body of evidence has shown that bilirubin has a protective effect on various oxidative stress-related diseases (9–15). We first reported a lower prevalence of micro- and macro-vascular complications in patients with diabetes and Gilbert syndrome, a congenital form of hyperbilirubinemia, implicating the protective role of bilirubin on diabetic vascular complications, including retinopathy (10). Low serum bilirubin levels have also been reported to be associated with an increased risk of diabetic retinopathy (16–18). However, because these studies are cross-sectional, it is still unclear how serum bilirubin levels are associated with diabetic retinopathy. In human, serum bilirubin levels are highly genetic (19, 20), while they are also influenced by environmental factors, including physiological and pathological conditions. Bilirubin functions as an antioxidant by reacting with ROS and its oxidized metabolites are excreted in urine (21). Lower serum bilirubin levels have been demonstrated in various chronic inflammatory and oxidative stress-related diseases, such as systemic lupus erythematosus, rheumatoid arthritis, and inflammatory bowel diseases (22–24). The explanation for these findings is likely to be the increased consumption of bilirubin. The oxidized metabolites of bilirubin in urine were designated as biopyrrins and reported to be increased in pathological oxidative stress-related conditions (25–27).

In this study, we therefore examined the association of the severity of retinopathy with serum bilirubin levels and various possible factors influencing bilirubin metabolism, including the oxidative stress marker, the inflammatory marker, urine biopyrrin levels and serum HO-1 levels.

## METHODS

### Study population

The study subjects consisted of 94 consecutive patients with diabetes mellitus admitted to Kyushu University Hospital from April 2011 to July 2012. Patients who suffered from hepatobiliary diseases with abnormal levels of aspartate aminotransferase, alanine aminotransferase or alkaline phosphatase and hemolytic anemia were excluded. All procedures were performed in accordance with the relevant guidelines and regulations. Informed consent was obtained from all participants. The study was approved by the ethics committee of Kyushu University Hospital.

### Clinical variables and definitions

In this study, venous blood samples were collected after overnight fasting. The serum bilirubin level was measured by the vanadate oxidation method. Total cholesterol, triglyceride and high-density lipoprotein cholesterol levels were measured using standard methods. The low-density lipoprotein (LDL) cholesterol level was calculated using the Friedewald formula. The hemoglobin A1c (HbA1c) value was determined using a standard high-performance liquid chromatography method. Throughout the paper, we present the National Glycohemoglobin Standardization Program (NGSP) value calculated as follows: JDC value + 0.4 (%)(18), and the International Federation of Clinical Chemistry and Laboratory Medicine (IFCC) mmol/mol units were converted using the NGSP converter for HbA1c, available at http://www.ngsp.org/convert1.asp. Serum high-sensitivity C-reactive protein (hs-CRP) was measured using an hs-CRP enzyme-linked immunosorbent assay (ELISA) kit (COSMO BIO Co., Ltd, Tokyo, Japan). We evaluated the oxidative stress state by measuring derivatives of reactive oxygen metabolites (d-ROMs). The d-ROM value was measured using the Free Carpe Diem program (Wismerll, Tokyo, Japan). The results of the d-ROM levels were expressed in an arbitrary unit called the Carratelli unit (U.CARR). Serum HO-1 concentration was determined by ELISA using a commercially available sandwich kit (Human HO-1 ELISA Kit, EKS-800, Stressgen Bioreagents/Assay Designs, Ann Arbor, MI, USA). For urinary biopyrrin measurement, the first urine in the morning was collected for measurement. All urine samples were immediately stored at −60°C and protected from light until measurement. Urinary biopyrrins were measured in duplicate, using a biopyrrin enzyme immunoassay kit based on a monoclonal antibody (Shino-test Co., Tokyo, Japan). The results were then corrected to the urinary concentration of creatinine, which was determined with the Creatinine Assay Kit (Cayman Chemical, Ann Arbor, MI). The urinary biopyrrin/creatinine ratio was used in subsequent analyses.

Retinopathy was assessed by a fundus examination by independent ophthalmologists. Classification was based on the modified Davis classification as follows: no diabetic retinopathy, simple diabetic retinopathy, pre-proliferative diabetic retinopathy, and proliferative diabetic retinopathy. Estimated glomerular filtration rate (eGFR) was calculated based on the Japanese Society of Nephrology CKD Practice Guide. Diabetic macroalbuminuria was defined as urinary albumin excretion ≥ 300 mg/g creatinine.

### Statistical analysis

Data were presented as mean ± standard deviation (SD) for normally distributed variables and as median (interquartile range) for nonparametric distribution. A two-sided P value of less than 0.05 was considered statistically significant. The patients were classified into three groups: the group with no retinopathy (NDR), the group with simple retinopathy (SDR), and the group with pre-proliferative or proliferative retinopathy (PDR). First, univariate analysis was performed to evaluate the relationship between the severity of retinopathy and various clinical factors. The three groups were compared using the analysis of variance test followed by the Dunnett post hoc analysis, Kruskal–Wallis test, and afterward by the Steel post hoc analysis or Pearson’s χ^2^ test. Next, multivariate logistic regression analysis was performed to evaluate the predictive value of serum bilirubin level for PDR. Because a significant difference was found among the three groups in serum bilirubin level, duration of diabetes and systolic blood pressure, we selected serum bilirubin levels alone for Model 1, serum bilirubin levels and duration of diabetes for Model 2, and serum bilirubin levels, duration of diabetes, systemic blood pressure, HbA1c and smoking were selected for Model 3.

Macroalbuminuria was strongly associated with PDR, and was thus excluded in this analysis. Other variables did not appreciably improve prediction and were not included in the model. Then, we plotted the ROC curve and calculated the AUC, sensitivity and specificity in each model. The optimal cut-off value of the bilirubin level for PDR was obtained from the Youden index [maximum = sensitivity + specificity − 1].

Second, to evaluate the significant determinants of serum bilirubin level, multiple regression analysis using variables thought to affect bilirubin metabolism was performed. All analyses were performed with JMP Version 14 statistical software (SAS Institute Inc., Cary, NC, USA).

## RESULTS

Table 1 shows the clinical characteristics of the participants. In univariate analysis, a significant difference was found among the NDR, SDR and PDR groups in systolic and diastolic blood pressure, duration of diabetes, serum total and indirect bilirubin levels, and macroalbuminuria. In post hoc analysis, compared with the NDR group, systolic and diastolic blood pressure were significantly (P<0.01) increased in the PDR group (P<0.01, Dunnett analysis), duration of diabetes was significantly increased in the SDR and PDR groups (P<0.05, P<0.01, respectively, Dunnett analysis), and serum bilirubin levels were significantly decreased in the PDR group (P<0.05, Steel analysis). Therefore, to evaluate the association between serum bilirubin levels and PDR, multivariate logistic regression analysis was performed. We selected serum bilirubin levels alone for Model 1, serum bilirubin levels and duration of diabetes for Model 2 and serum bilirubin levels, duration of diabetes, systemic blood pressure, HbA1c and smoking were selected for Model 3. As shown in Table 2, serum total bilirubin levels alone were significant determinants for PDR (OR 0.770, 95% CI 0.639–0.929, P=0.006). Model 2 showed that both serum bilirubin levels and disease duration were significant determinants for PDR (OR 0.772, 95% CI 0.622–0.959, P=0.020 and OR 1.087, 95% CI 1.031–1.146, P=0.002, respectively). Model 3 showed that serum bilirubin levels, disease duration and systolic blood pressure were significant determinants for PDR (OR 0.737, 95% CI 0.570–0.952, P=0.020; OR 1.085, 95% CI 1.024–1.149, P=0.006; OR 1.036, 95% CI 1.011–1.062, P=0.005; OR 1.242, respectively), but not HbA1c or smoking. ROC analysis was subsequently performed to evaluate the relative ability of each model to predict PDR. As shown in Fig 1, the area under the curve (AUC) in Model 1 was 0.677 (specificity=59.4%, sensitivity=70.0%). AUC in Model 2 was 0.780 (specificity=86.4%, sensitivity=63.0%). The AUC in Model 3 was 0.832 (specificity=71.2%, sensitivity=81.5%). Serum bilirubin level alone was a significant determinant for PDR, and its predictive ability was further increased by the addition of duration of diabetes and systolic blood pressure.

**Table 1.** Characteristics of total patients, patients without retinopathy, patients with simple retinopathy, and patients with pre-proliferative or proliferative retinopathy.

|  | Total | NDR | SDR | PDR | P Value |
| --- | --- | --- | --- | --- | --- |
| N | 94 | 49 | 15 | 30 |  |
| Age (years) | 58.6 ± 15.0 | 57.8 ± 16.3 | 66.1 ± 11.0 | 56.0 ± 13.6 | 0.089† |
| Gender (male) | 53 (56.4%) | 30 (61.2%) | 6 (40.0%) | 17 (56.7%) | 0.349‡ |
| Systolic blood pressure (mmHg) | 135.5 ± 24.4 | 126.4 ± 19.1 | 140.7 ± 23.0 | 147.8 ± 27.3** | <.001† |
| Diastolic blood pressure (mmHg) | 78.7 ± 11.9 | 75.1 ± 10.4 | 78.9 ± 10.1 | 84.4 ± 12.9** | 0.002† |
| Body mass index (kg/m<sup>2</sup>) | 24.8 ± 5.0 | 24.5 ± 5.9 | 24.8 ± 5.2 | 24.7 ± 5.5 | 0.785† |
| Duration of diabetes (years) | 7.0 (2.0-15.5) | 3.0 (1.0-9.5) | 9.0 (6.5-17.0)* | 12.0 (9.0-21.0)** | <.001§ |
| Smoking | 48 (51.1%) | 25 (51.0%) | 7 (46.7%) | 16 (53.3%) | 0.915‡ |
| HbA1c (%) | 9.3 ± 2.0 | 9.4 ± 2.1 | 9.2 ± 2.0 | 9.4 ± 2.0 | 0.573† |
| Serum total bilirubin (mg/dl) | 0.75 (0.60-0.90) | 0.80 (0.60-1.00) | 0.80 (0.80-0.90) | 0.65 (0.50-0.80)* | 0.014§ |
| Serum indirect bilirubin (mg/dl) | 0.60 (0.50-0.70) | 0.60 (0.50-0.78) | 0.70 (0.55-0.70) | 0.50 (0.40-0.60)* | 0.031§ |
| LDL cholesterol (mg/dl) | 117.8 ± 45.6 | 113.0 ± 37.5 | 114.3 ± 38.8 | 127.4 ± 58.7 | 0.381† |
| HDL cholesterol (mg/dl) | 47.7 ± 14.5 | 45.7 ± 12.5 | 45.1 ± 11.4 | 52.1 ± 17.9 | 0.125† |
| Triglycerides (mg/dl) | 125.5 (79.8-171.3) | 131.0 (94.0-171.5) | 123.0 (82.0-193.0) | 129.5 (71.5-154.3) | 0.841§ |
| eGFR (ml/min/1.73m<sup>2</sup>) | 80.3 (65.1-98.4) | 84.9 (66.8-102.1) | 74.0 (66.1-86.2) | 69.9 (35.9-108.6) | 0.215§ |
| Macroalbuminuria | 14 (15.1%) | 1 (2.1%) | 3 (20.0%) | 10 (33.3%) | <.001‡ |
| Treatment |  |  |  |  |  |
| Diet only | 4 (4.3%) | 1 (2.0%) | 0 (0.0%) | 3 (10.0%) | 0.158‡ |
| Oral hypoglycemic agent | 27 (28.7%) | 16 (32.7%) | 5 (33.3%) | 6 (20.0%) | 0.440‡ |
| Insulin | 58 (61.7%) | 29 (59.2%) | 9 (60%) | 20 (66.7%) | 0.793‡ |
| GLP1 analogue | 5 (5.3%) | 3 (6.1%) | 1 (6.7%) | 1 (3.3%) | 0.839‡ |
| hs-CRP (ng/ml) | 606 (331-1388) | 499 (305-1405) | 948 (205-1880) | 653 (421-1330) | 0.704§ |
| d-ROMs | 295 (262-364) | 289 (261-364) | 319 (265-343) | 296 (258-387) | 0.728§ |
| Serum hemeoxygenase-1 (ng/ml) | 5.85 (3.90-7.39) | 6.00 (3.89-7.84) | 6.68 (4.18-8.15) | 5.36 (3.49-6.82) | 0.299§ |
| Urinary biopyrrins (U/g/cre) | 1.20 (0.68-2.34) | 1.21 (0.68-2.21) | 1.22 (0.98-3.43) | 1.05 (0.61-2.12) | 0.175§ |

NDR diabetic patients without retinopathy; SDR diabetic patients with simple retinopathy; PDR diabetic patients with pre-proliferative or proliferative retinopathy.
† Calculated using the Analysis of variance test, followed by Dunnett analysis
‡ Calculated using the Pearson's $\chi^2$ test.
§ Calculated using the Kruskal-Wallis test, followed by Steel analysis
\* P<0.05 vs. NDR; \*\* P<0.01 vs. NDR

**Table 2.** Multivariate logistic regression models predicting PDR (pre-proliferative or proliferative retinopathy)

|  | Model 1 | Model 2 | Model 3 |
| --- | --- | --- | --- |
|  | OR (95%CI) | OR (95%CI) | OR (95%CI) |
|  | P Value | P Value | P Value |
| Serum bilirubin (10×)<br>(mg/dl) | 0.770 (0.639-0.929)<br>0.006 | 0.772 (0.622-0.959)<br>0.020 | 0.737 (0.570-0.952)<br>0.020 |
| Duration of diabetes<br>(year) |  | 1.087 (1.031-1.146)<br>0.0020 | 1.085 (1.024-1.149)<br>0.006 |
| Systolic blood pressure<br>(mmHg) |  |  | 1.036 (1.011-1.062)<br>0.005 |
| HbA1c<br>(%) |  |  | 1.242 (0.909-1.698)<br>0.174 |
| Smoking |  |  | 0.981 (0.322-2.987)<br>0.973 |

Data are represented as odds ratio (OR) and 95%CI (confidence interval). Model 1: serum bilirubin levels alone; Model 2: serum bilirubin levels and duration of diabetes; Model 3: serum bilirubin levels, duration of diabetes, systolic blood pressure, HbA1c and smoking

**Figure 1.**
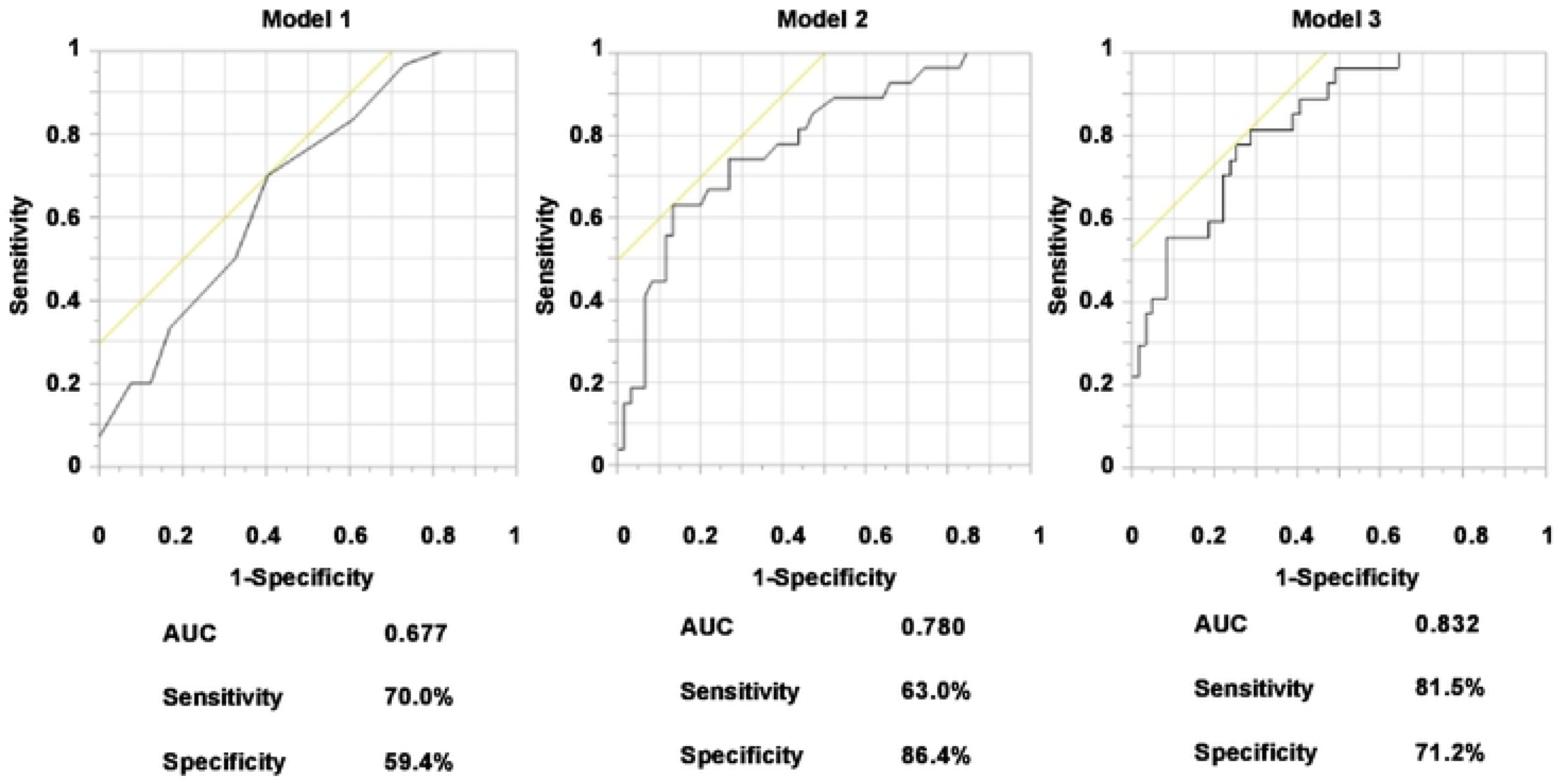
AUC of various models to predict PDR (pre-proliferative retinopathy or proliferative retinopathy) evaluated by ROC curves. ROC curves demonstrate the relative ability of serum bilirubin levels (Model 1), serum bilirubin levels and duration of diabetes (Model 2) and serum bilirubin levels, duration of diabetes, systolic blood pressure, HbA1c and smoking (Model 3) to predict PDR. AUC, area under the curve. ROC, receiver operating characteristic.

In this study, multiple regression analysis using variables thought to affect bilirubin metabolism was performed to evaluate the significant determinants for serum bilirubin levels. This analysis showed that serum bilirubin levels were negatively associated with serum hs-CRP levels and PDR (P=0.018, P=0.024, respectively), and positively associated with urinary biopyrrin levels (P=0.014) (Table 3).

**Table 3.** Evaluation of significant determinants for serum bilirubin levels by multivariate regression analysis

| Variables | Regression coefficient | 95% CI (lower upper) |  | P Value |
| --- | --- | --- | --- | --- |
| Systolic blood pressure | 0.0157 | -0.010 | 0.041 | 0.225 |
| Age | 0.0370 | -0.008 | 0.082 | 0.104 |
| Gender (male) | -0.1144 | -1.322 | 1.093 | 0.851 |
| Body mass index | 0.0022 | -0.119 | 0.123 | 0.971 |
| Duration of diabetes | 0.0237 | -0.044 | 0.091 | 0.487 |
| Smoking | -0.3420 | -1.581 | 0.897 | 0.584 |
| HbA1c | 0.2392 | -0.037 | 0.516 | 0.089 |
| eGFR | 0.0198 | -0.001 | 0.041 | 0.067 |
| hsCRP | -0.0002 | -0.000 | 0.000 | 0.018* |
| Urinary biopyrrins | 0.3040 | 0.064 | 0.544 | 0.014* |
| Hemeoxygenase-1 | 0.1178 | -0.110 | 0.346 | 0.306 |
| dROMs | -0.0035 | -0.010 | 0.003 | 0.303 |
| PDR | -1.6360 | -3.051 | -0.221 | 0.024* |

Data are presented as partial regression coefficient and 95% CI.

## DISCUSSION

We first reported a lower prevalence of retinopathy in patients with diabetes and Gilbert syndrome, a congenital form of hyperbilirubinemia, indicating the protective role of bilirubin (8). Many reports have also shown that serum bilirubin levels are negatively associated with a risk of diabetic retinopathy (16–18). In this study, we found that serum bilirubin levels alone were significant determinants for PDR (pre-proliferative or proliferative retinopathy), and its predictive ability was further increased by the addition of duration of diabetes and systolic blood pressure. Because serum bilirubin is a strong endogenous antioxidant, it is most likely that low antioxidant activity due to low serum bilirubin levels may cause the worsening of retinopathy because oxidative stress has been thought to play an important role in the pathogenesis of diabetic retinopathy (4–7). However, it is also possible that low serum bilirubin levels may be a result of PDR or other coexisting diabetic complications. Bilirubin functions as an antioxidant in vivo by reacting with ROS and then being consumed. Serum bilirubin levels may be decreased in diabetic patients with PDR who are in increased oxidative conditions. In the present study, we measured urinary biopyrrin levels to evaluate the possibility of the increased consumption of serum bilirubin. Unexpectedly, the present study showed that urinary biopyrrin levels were positively correlated with serum bilirubin levels, suggesting that decreased serum bilirubin levels in patients with PDR may be due to decreased production of bilirubin rather than increased consumption.

Bilirubin is mainly generated from heme degradation by the ubiquitously expressed HO-1, the rate-limiting enzyme involved in heme catabolism. Therefore, the present results suggested that the HO-1 expression/activity may be impaired in patients with PDR, and thus serum bilirubin production was decreased. HO-1 is induced strongly by numerous stress stimuli such as ROS, pro-inflammatory cytokines and heme itself (28). This HO-1 response is thought to play an important role in counteracting increased oxidative stress and inflammation and thus maintaining redox homeostasis. In the diabetic state, HO-1 expression was reported to increase in various tissues, such as kidney, circulating monocytes and lymphocytes, probably in response to increased oxidative stress and chronic inflammation (29–31). In contrast, several reports have shown inconsistent findings that HO-1 expression was decreased in some tissues of diabetic patients and animal models (32–35). One possible explanation for such inconsistency might be the variation in HO-1 expression at different stages of diabetes. In the retina, one report showed increased expression of HO-1 in short-term diabetes (31). Another interesting report showed that HO-1 expression in the retina was increased in 8-week-old db/db mice, while it was decreased in 20-week-old db/db mice, indicating that the HO-1 response might be impaired in late-stage diabetes (36). Indeed, diminished HO-1 mRNA expression in retinal pigment epithelial cells obtained from diabetic donors was also reported (32). However, the mechanism underlying impaired HO-1 response in late-stage diabetes is unknown. In recent years, nuclear factor erythroid 2-related factor 2 (Nrf2) has attracted increasing attention as a major transcriptional regulator of antioxidant proteins, including HO-1. Of great interest, increasing evidence has indicated that the Nrf2/antioxidant response might be impaired across a wide range of tissue types in diabetes (37), although the mechanisms underlying Nfr2 dysfunction also remain to be elucidated. HO-1 levels in the retina of STZ-induced diabetic rats were reported to reach their peak at 4 weeks after onset of diabetes, then decrease at 6 weeks, combined with accordant changes of Nrf2 expression (38). Furthermore, overexpression of HO-1 by hemin protected the development retinopathy at 8 weeks (38). Collectively, all these findings indicated that decreased serum bilirubin levels in patients with PDR might be associated with the impaired Nrf2/HO-1 pathway response. However, to confirm this hypothesis, the relationship between diabetic retinopathy, serum bilirubin levels and the Nrf2/HO-1 pathway should be evaluated in long-term prospective studies.

This study has several limitations. First, the sample size was small and the study was cross-sectional. In this small-scale study, we found that serum bilirubin levels were significantly associated with PDR, but not SDR. Larger scale studies are needed to evaluate in detail the association between the severity of retinopathy and serum bilirubin levels. Second, we measured serum HO-1 levels in this study. Elevated serum HO-1 levels were reported in individuals with newly diagnosed type 2 diabetes compared with control individuals probably in response to increased oxidative stress (39). In this study, serum HO-1 levels were not significantly different in the PDR group compared with the NDR group, but they tended to be decreased despite decreased bilirubin levels in PDR. This finding might support the impaired HO-1 response in PDR. However, the relationship between serum bilirubin levels, HO-1 levels in serum and in the tissues including the retina is not fully understood. This should be also evaluated in future studies.

In conclusion, PDR was negatively associated with serum bilirubin levels in patients with diabetes. This negative association may be due to a decreased production of bilirubin rather than its increased consumption, considering the positive correlation between serum bilirubin levels and urinary biopyrrin levels, which are oxidized metabolites of bilirubin.

## ACKNOWLEDGMENTS

This work was supported in part by a grant for the Creation of Innovation Centers for Advanced Interdisciplinary Research Areas Program from the Ministry of Education, Culture, Sports, Science and Technology of Japan (Funding program “Innovation Center for Medical Redox Navigation”).

